# RUNX1 mitotically bookmarks target genes that are important for the mammary epithelial-to-mesenchymal transition

**DOI:** 10.1101/511410

**Authors:** Joshua T. Rose, Eliana Moskovitz, Joseph R. Boyd, Jonathan A. Gordon, Nicole A. Bouffard, Andrew J. Fritz, Anuradha Illendula, John H. Bushweller, Jane B. Lian, Janet L. Stein, Gary S. Stein, Sayyed K. Zaidi

**Affiliations:** Department of Biochemistry and University of Vermont Cancer Center, Robert Larner College of Medicine, 89 Beaumont Avenue, Burlington, VT 05405, USA; Microscopy Imaging Center at the Robert Larner College of Medicine, 89 Beaumont Avenue, Burlington, VT 05405, USA; Department of Molecular Physiology and Biological Physics, University of Virginia, Charlottesville, VA 22908, USA

**Keywords:** Mammary phenotype, epithelial phenotype, RUNX1, mitotic gene bookmarking

## Abstract

RUNX1 has recently been shown to play an important role in determination of mammary epithelial cell identity. However, mechanisms by which loss of the RUNX1 transcription factor in mammary epithelial cells leads to epithelial-to-mesenchymal transition (EMT) are not known. Here, we report mitotic bookmarking of genes by RUNX1 as a potential mechanism to convey regulatory information through successive cell divisions for coordinate control of mammary cell proliferation, growth, and identity. Genome-wide RUNX1 occupancy profiles for asynchronous, mitotically enriched, and early G1 breast epithelial cells reveal RUNX1 is retained during the mitosis to G1 transition on protein coding and long non-coding RNA genes critical for mammary epithelial proliferation, growth, and phenotype maintenance. Disruption of RUNX1 DNA binding and association with mitotic chromosomes alters cell morphology, global protein synthesis, and phenotype-related gene expression. Together, these findings show for the first time that RUNX1 bookmarks a subset of epithelial-related genes during mitosis that remain occupied as cells enter the next cell cycle. Compromising RUNX1 DNA binding initiates EMT, an essential first step in the onset of breast cancer.

**Significance:** This study elucidates mitotic gene bookmarking as a potential epigenetic mechanism that impacts breast epithelial cell growth and phenotype and has potential implications in breast cancer onset.

## INTRODUCTION

Breast cancer arises from a series of acquired mutations and epigenetic changes that disrupt normal mammary epithelial homeostasis and create multi-potent cells that can differentiate into biologically unique and clinically distinct subtypes (1–6). Epithelial-to-mesenchymal transition (EMT)—a trans-differentiation process through which mammary epithelial cells acquire the aggressive mesenchymal phenotype—is a key driver of breast cancer progression, invasion and metastasis (7–12). Transcription factors Snail, Slug, Twist, and Zeb1/2 contribute to EMT during early, normal development and have also been implicated in invasion (13–16). Despite accumulating evidence that defines a broad understanding of EMT regulation and maintenance of the epithelial phenotype (7–12), the mechanism(s) by which mammary cells maintain their epithelial phenotype is unknown.

Runt-Related Transcription Factor 1 (RUNX1/AML1) is required for hematopoietic lineage specification during development and hematopoiesis throughout life (17–30). In addition to the recognized role in hematological malignancies, RUNX1 has been recently identified as a key player in breast cancer development and tumor progression (31–38). Findings from our group (39), reinforced by studies from others (40, 41), have shown that RUNX1 plays a critical role in maintaining breast epithelial phenotype and prevents EMT through transcriptional regulation of genes (e.g., the EMT marker and a key cell adhesion protein E-cadherin) involved in fundamental cellular pathways. However, mechanisms by which RUNX1 prevents EMT have not been identified.

Mitotic gene bookmarking, i.e. transcription factor binding to target genes during mitosis for transcriptional regulation following cell division, is a key epigenetic mechanism to convey and sustain regulatory information for cell proliferation, growth, and cell identity from parent to progeny cells (42–49). We have established that RUNX proteins, as well as other phenotypic transcription factors that include MYOD and CEBPα, mitotically bookmark RNA Pol I- and II-transcribed genes in osteoblasts and leukemia cells for coordinate control of cell growth, proliferation and phenotype (50–57). It is increasingly evident that mitotic gene bookmarking by transcription factors is a key mechanism to determine and maintain cell fate across successive cell divisions (58–70).

We addressed the hypothesis that RUNX1 maintains the breast epithelial phenotype by mitotic bookmarking of genes that support mammary epithelial proliferation, growth, and phenotype for expression in the next cell cycle. Fluorescence confocal microscopy of fixed and live mammary epithelial cells revealed that RUNX1 is present on chromosomes throughout mitosis and colocalizes with upstream binding transcription factor (UBF), a subunit of RNA Pol I transcriptional machinery (71). To identify genes occupied by RUNX1, we performed chromatin immunoprecipitation coupled with high throughput sequencing (ChIP-Seq) using a RUNX1-specific antibody on mitotic, G1, and asynchronous normal mammary epithelial MCF10A cells. As expected, ribosomal RNA genes, regulated by the RNA Pol I transcriptional machinery, were occupied by RUNX1. A fluorescence-based, global protein synthesis assay showed reduced protein synthesis when RUNX1 DNA binding was perturbed using a small molecule inhibitor. Importantly, ChIP-Seq revealed that, in mitosis, RUNX1 associates with RNA Pol II regulated genes specifically involved in maintenance of the epithelial phenotype and EMT progression. Strikingly, disruption of RUNX1 DNA binding, which is required for association with mitotic chromosomes (56), results in loss of the epithelial phenotype and acquisition of mesenchymal properties that are accompanied by changes in expression of associated genes and pathways and represent early events in the onset of breast cancer. These findings implicate RUNX1 occupancy of target genes at the mitosis into G1 transition in regulating the normal breast epithelial phenotype.

## RESULTS

### RUNX1 associates with mitotic chromatin and occupies target genes

To investigate subcellular localization of RUNX1 in normal mammary epithelial cells, we performed immunofluorescence (IF) microscopy in actively proliferating MCF10A cells and imaged cells undergoing spontaneous mitoses. We observed that RUNX1 is localized on mitotic chromatin at all topologically identified substages of mitosis (Fig. 1; top panels). Two distinct types of foci are detectable on mitotic chromosomes: 2-8 large punctate foci that appear to be allelic as well as numerous smaller foci that are distributed across the chromosomes (Fig. 1; bottom panels, white arrowheads). In all replicates, important secondary-antibody-only controls were included to confirm specificity of RUNX1 signal on mitotic chromosomes (Suppl. Fig. 1). In the interphase nuclei, RUNX1 exhibited the characteristic punctate distribution (Fig. 1; interphase panel). To ensure reproducibility of our findings, the IF experiments were repeated at least 3 times and, at the minimum, 20 interphase and mitotic cells were imaged.

**Figure 1.**
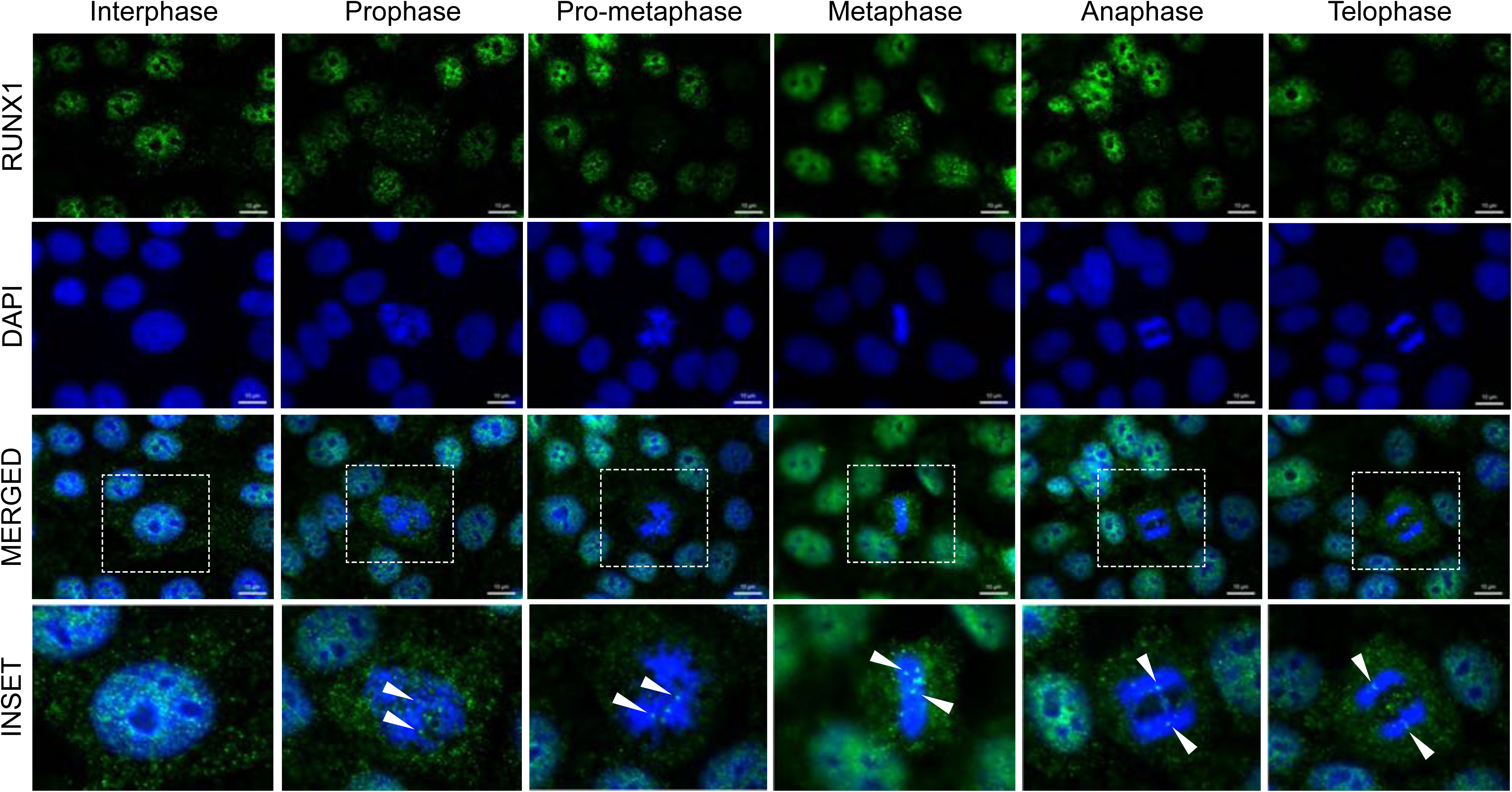
RUNX1 associates with DNA during interphase and remains bound throughout mitosis in the form of major and minor foci. Representative immunofluorescent images of interphase and mitotic MCF10A breast epithelial cells showing subcellular localization of RUNX1, identified using a specific antibody, throughout mitosis. Mitotic cells were further classified into substages of mitosis based on DAPI topology. RUNX1 – Green (top row), DAPI – Blue (second row from top). Merged channel images (third row from top) contain an outlined region magnified in the bottom row labeled “inset”. White arrows highlight major Runx1 foci on mitotic chromatin. Three independent biological replicates were performed, and at least 20 cells for each mitotic substage were analyzed.

Multiple reports have indicated that formaldehyde fixation can prevent regulatory protein detection on mitotic chromosomes (58, 69). To further confirm that association of RUNX1 with mitotic chromosomes is not under-represented because of formaldehyde fixation, we examined the localization of RUNX1-EGFP in actively proliferating, unfixed MCF10A cells. Consistent with our findings in fixed and synchronized cells, RUNX1-EGFP was associated with chromosomes in live MCF10A cells undergoing mitosis (Suppl. Fig. 2.). Together, these findings establish that RUNX1 foci are present on chromosomes at all stages of mitosis under physiological conditions in actively dividing, unfixed breast epithelial cells and, in agreement with our previous findings, are equally distributed into resulting progeny cells (57).

To experimentally address whether RUNX1 presence on mitotic chromosomes reflects occupancy of target genes, MCF10A cells were synchronized in mitosis using nocodazole (50ng/mL; Supp. Fig. 3A). Nocodazole dose and treatment were empirically determined to minimize toxic effects of the drug, while maximizing mitotic enrichment. Mitotic cells were collected by mitotic shake-off and purity of harvested cells was confirmed by the presence of H3pS28 in >70% of cells. We chose the H3pS28 mark to identify mitotic cells because this histone mark is highly specific to condensed chromosomes during mitosis; more commonly used H3pS10 mark is additionally observed in early G1 cells and has also been associated with replicating centers in S-phase (72, 73). A parallel, nocodazole-treated cell population was released into early G1 by replacing nocodazole-containing growth medium with fresh, nocodazole-free, growth medium and was harvested 3 hours post-release (Suppl. Fig. 3B). Western blot analysis of whole cell lysates from the three cell populations showed expected levels of expression for cell cycle-specific proteins Cyclin B and CDT1 (Suppl. Fig. 3C). FACS profiles of the cell populations confirmed the characteristic enrichment of blocked cells in mitosis (Supp. Fig. 3D; Mitotic) and release into G1 upon media replacement (Supp. Fig. 3D; G1) when compared to asynchronous cells (Supp. Fig. 3D; Asynch). To determine whether RUNX1 remains bound to target genes during mitosis, ChIP-Seq was performed on Asynch, Mitotic, and G1 MCF10A cells using a RUNX1 specific antibody (Fig. 2A). Enriched regions (peaks called by MACS2) were compared using k-means clustering (k=4) of normalized enrichment profiles of the three cell populations. This analysis revealed subsets of genes that were either shared across the three groups or were specific for each, indicating dynamic binding of RUNX1 during and immediately after mitosis (Fig. 2A). Peak calling identified RUNX1 occupancy of both protein coding and long non-coding RNA (lncRNA) genes. Specifically, RUNX1 occupied 1070 genes in cell population not in G1 or M phase (Fig. 2A; green bar) and 1095 genes in G1-enriched cells (Fig. 2A; light red bar). Importantly, RUNX1 occupied 551 genes (413 protein coding and 138 lncRNAs) in mitotically enriched MCF10A cells (Fig. 2A; blue bar).

**Figure 2.**
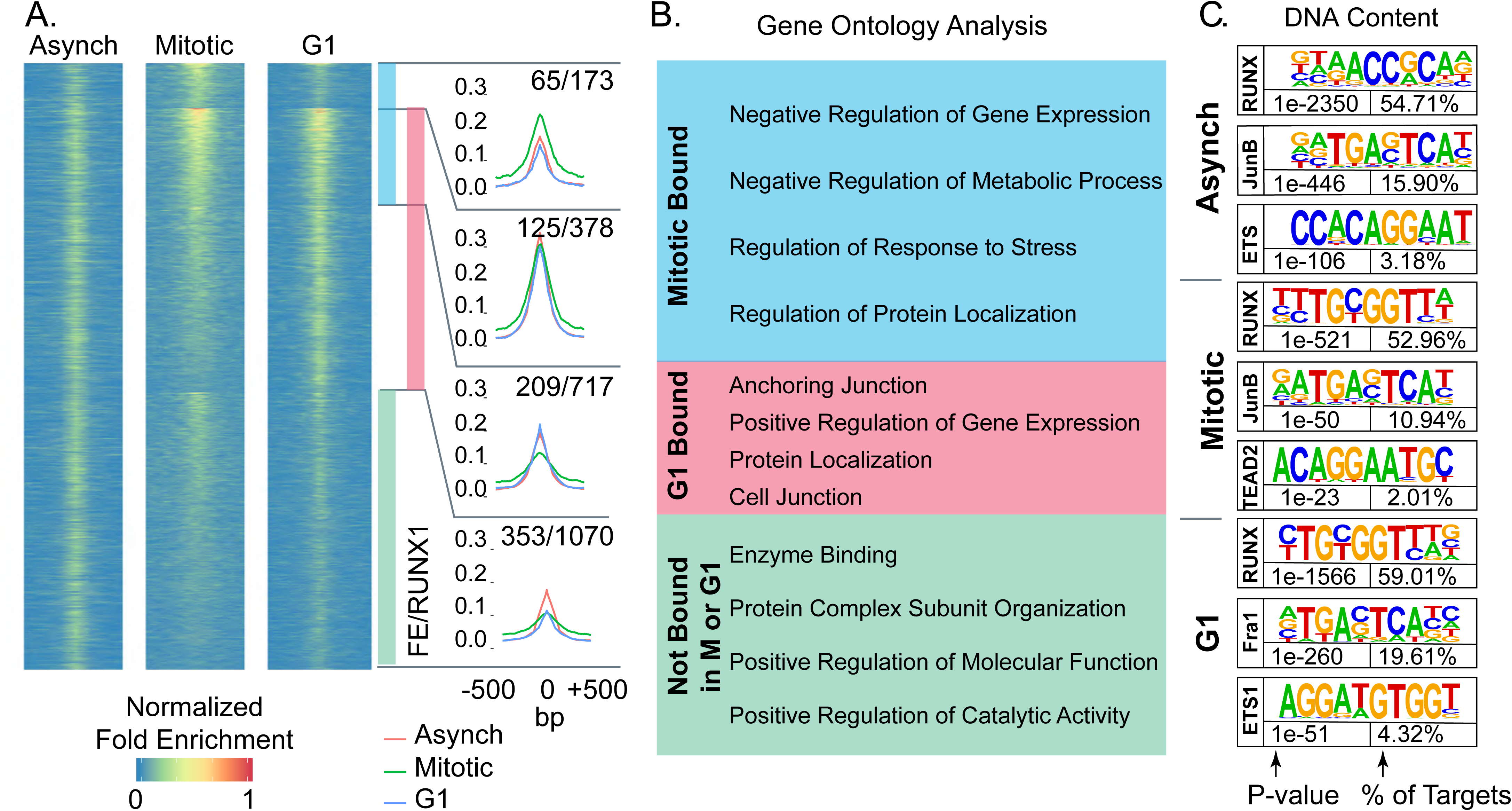
RUNX1 occupies protein coding genes and long non-coding RNAs across asynchronous, mitotic, and G1 populations of MCF10A breast epithelial cells. A) Heatmaps showing peaks called between A, M, and G1 MCF10A cells (left, middle, and right respectively). Cumulative occupancy of RUNX1 is shown as line graphs and genes that occupied by RUNX1 in each of the three cell populations are shown. Shown also are genes that are deregulated upon RUNX1 downregulation. For example, 1070 genes are bound by RUNX1 in cells that are neither in M nor G1 cells (green bar graph) and of those, 353 genes are deregulation upon RUNX1 deregulation. B) Gene ontology analysis of RUNX1-bound genes identifies key regulatory pathways that are similar or unique in the three cell populations. C) Motif analysis of A, M, and G1 MCF10A cells reveals RUNX motif as one of the top motifs in all cell populations. Binding sites for key transcription factors that are known to cooperate with RUNX1 are also identified.

Functional relevance of RUNX1 occupancy in the three cell populations was determined by comparing RUNX1-occupied genes with those that are differentially regulated upon shRNA-mediated RUNX1 knockdown (39). Of the 1070 genes occupied by RUNX1 in cell populations not in G1 or M phases, 353 genes were deregulated upon RUNX1 depletion (Fig. 2A). Importantly, RUNX1 depletion deregulated 399 of 1268 RUNX1-bound genes in the M and G1 populations. These findings reveal that several hundred target genes are bookmarked by RUNX1 during mitosis and transcriptionally regulated in normal mammary epithelial cells.

To identify cellular processes and pathways that are comprised of RUNX1-bookmarked genes, we performed gene set enrichment analysis (GSEA) on genes bound by RUNX1 during mitosis or G1, or not bound in either cell cycle stage (Fig. 2B). Interestingly, most genes bookmarked by RUNX1 during mitosis were associated with negative regulation of gene expression and metabolic process (Fig. 2B; blue box). Consistent with a cellular requirement to reattach and enter the next cell cycle and fully resume transcription, genes bound during G1 were primarily enriched in biological processes involving cell anchorage, protein localization and positive regulation of gene expression (Fig. 2B; red box). ChIP-seq results were further validated by motif analysis of RUNX1-bound peaks, which showed that the RUNX motif was the most enriched motif in all three cell populations (Fig. 2C). Importantly, RUNX1-bound genomic regions were also enriched in motifs for transcription factors (e.g., Fra1, JunB, ETS) known to cooperate with RUNX1 for gene regulation (74)(Fig. 2C). Together, these findings indicate that RUNX1 occupies genes involved in cell proliferation, growth, and phenotype during mitosis in normal mammary epithelial cells.

### RUNX1 mitotically bookmarks RNA Pol I-transcribed genes that control cell growth

Our ChIP-Seq results revealed that RUNX1 occupies rDNA repeats in MCF10A mammary epithelial cells; all three MCF10A cell populations (Asynch, Mitotic and G1) exhibited significant fold enrichment within the promoter region of hrDNA (Fig. 3A), suggesting a potential role for RUNX1 in regulating rRNA genes in MCF10A cells. We confirmed this finding in actively proliferating MCF10A cells by immunofluorescence microscopy for antibodies specific against RUNX1 and upstream binding factor (UBF), a transcriptional activator that remains bound to rRNA genes during mitosis (75). We observed large RUNX1 foci colocalizing with UBF throughout each stage of mitosis (Fig 3B and Suppl. Fig. 4). Colocalization between RUNX1 and UBF was validated by confocal microscopy. Line scans of MCF10A cells show that although RUNX1 and UBF occupy distinct nuclear microenvironments in interphase (n=15), both proteins substantially colocalize in metaphase (n=15)(Suppl. Fig. 4). Taken together, these findings establish RUNX1 binding to ribosomal DNA repeat regions by ChIP-Seq (Fig. 3A) and confirmed at the cellular level by confocal microscopy (Fig 3B & Suppl. Fig. 4).

**Figure 3.**
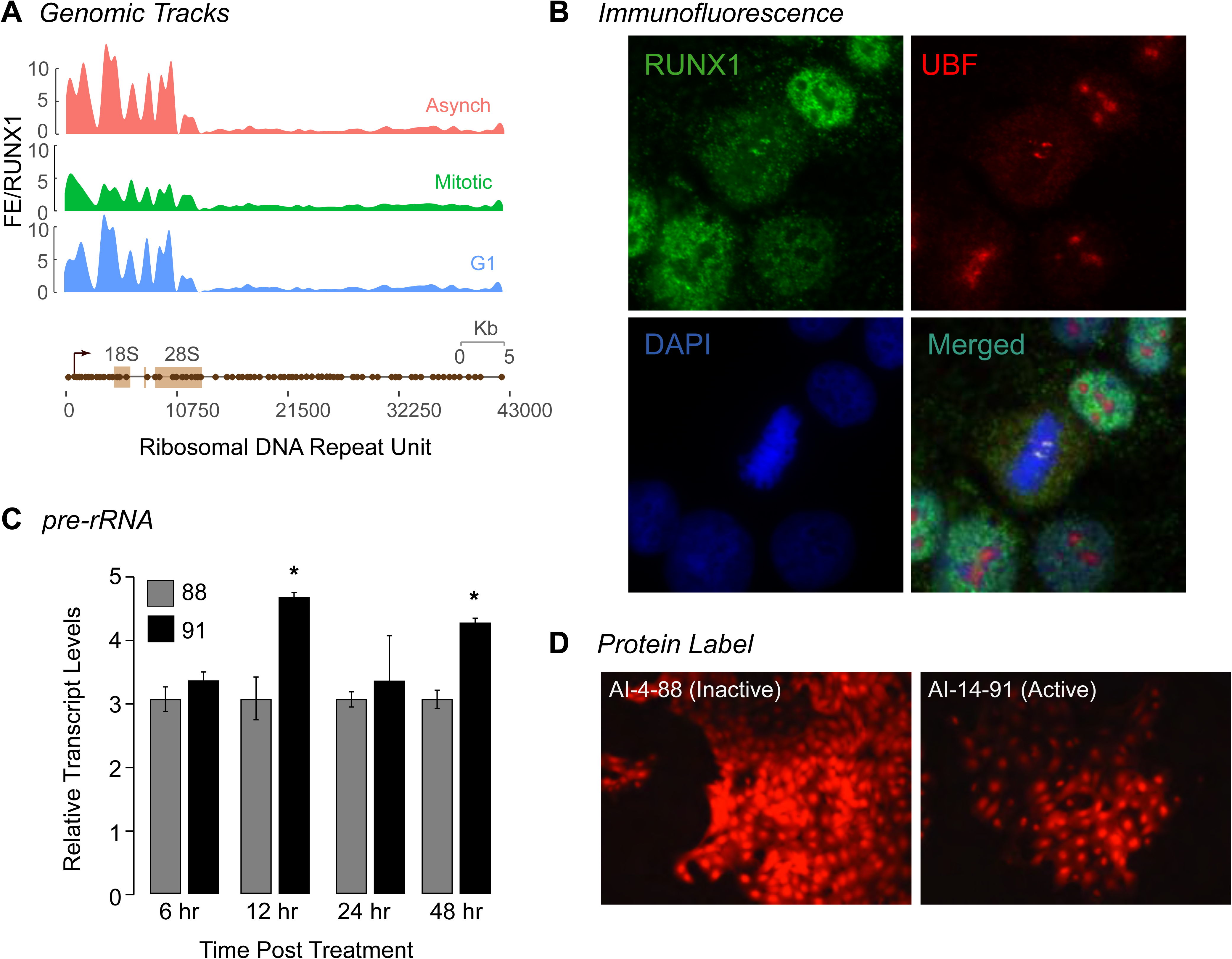
RUNX1 bookmarks rDNA promoter repeat regions and affects both pre-rRNA and global protein expression. A) ChIP-Seq tracks of A, M, and G1 (top, middle, bottom respectively) MCF10A cells mapped against rDNA repeat regions. B) A representative metaphase MCF10A cell, stained for RUNX1 (green) and UBF1 (red) localization, is shown demonstrating that the two proteins colocalize during mitosis (merged). Cells are also counter stained with DAPI to visualize DNA (blue) and identify mitosis substages. C) qRT-PCR data of pre-rRNA in actively proliferating MCF10A cells treated with either active (AI-14-91) or inactive (AI-4-88) compounds for 6, 12, 24, or 48hrs. Expression of pre-rRNA was normalized relative to Beta Actin expression. Graph represents three independent biological replicates. Asterisks represents a *p* value of <0.05. D) Representative fluorescence microscopy images of global protein synthesis from MCF10A cells treated with either AI-4-88 (left) or AI-14-91 (right) for 24hr at 20μM (n=2). Intensity of red fluorescence at 580nm emission indicates nascent protein synthesis. All images were taken with 1000ms exposures.

We experimentally addressed the hypothesis that RUNX1 regulates ribosomal RNA gene expression by using a pharmacological inhibitor of RUNX1. The small molecule inhibitor AI-14-91, which has been extensively characterized by several groups, interferes with RUNX1-CBFβ interaction and disrupts RUNX1 DNA binding (76, 77). We examined the effect of RUNX1 inhibitor on pre-rRNA expression and found that pre-rRNA expression was significantly increased at 12hr and 48hr time points after treatment of asynchronous cells with the specific RUNX1 inhibitor but not the control inactive compound AI-4-88, indicating that RUNX1 suppresses rRNA gene expression in normal mammary epithelial cells (Fig. 3C). Because levels of rRNA directly correlate with global protein synthesis, a fluorescent-based detection method was used to measure newly synthesized proteins. Cells treated with AI-14-91 for 24hr or 48hr showed a moderate change in levels of global protein synthesis in comparison to AI-4-88 control-treated cells under identical conditions (n=3; Fig. 3D). Together, our results demonstrate that RUNX1 bookmarks RNA Pol I regulated rRNA genes during mitosis and transcriptionally represses them during interphase with moderate impact on global protein synthesis in normal mammary epithelial cells.

### RUNX1 occupies RNA Pol II-transcribed genes involved in hormone-responsiveness and cell phenotype during mitosis

Using RUNX1-bookmarked genes, gene set enrichment analysis (GSEA) was performed to identify regulatory pathways (Fig. 4A). In agreement with known roles of RUNX1 (78–82), the top 10 pathways identified were those involved in regulation of G2M Checkpoint, E2F targets, p53, and DNA repair (Fig. 4A). Consistent with our finding that RUNX1 bookmarks and regulates rRNA genes, one of the pathways identified is mTOR signaling, a pathway that is required for cell growth and is a therapeutic target in breast cancers (83, 84). Relevant to the normal mammary epithelial phenotype, both early and late estrogen responsive gene sets significantly overlap with RUNX1 mitotically bookmarked genes (Fig. 4A). Because estrogen plays vital roles in promoting proliferative phenotypes of mammary epithelial cells (85–87), we interrogated RUNX1 bookmarked genes to identify those bound by RUNX1 and ERα in MCF7 cells, where RUNX1 contributes to higher order genome organization (Fig. 4B) (88, 89). Using publicly available datasets of ERα genome-wide occupancy and estradiol-regulated gene expression (GSE40129)(90), we find that a subset of genes mitotically bookmarked by RUNX1 is also bound by ERα, and either up or down regulated in response to estradiol. These findings indicate that RUNX1-bookmarked genes are involved in pathways that control hormone-responsiveness, proliferation and growth of normal mammary epithelial cells (Fig. 4B).

**Figure 4.**
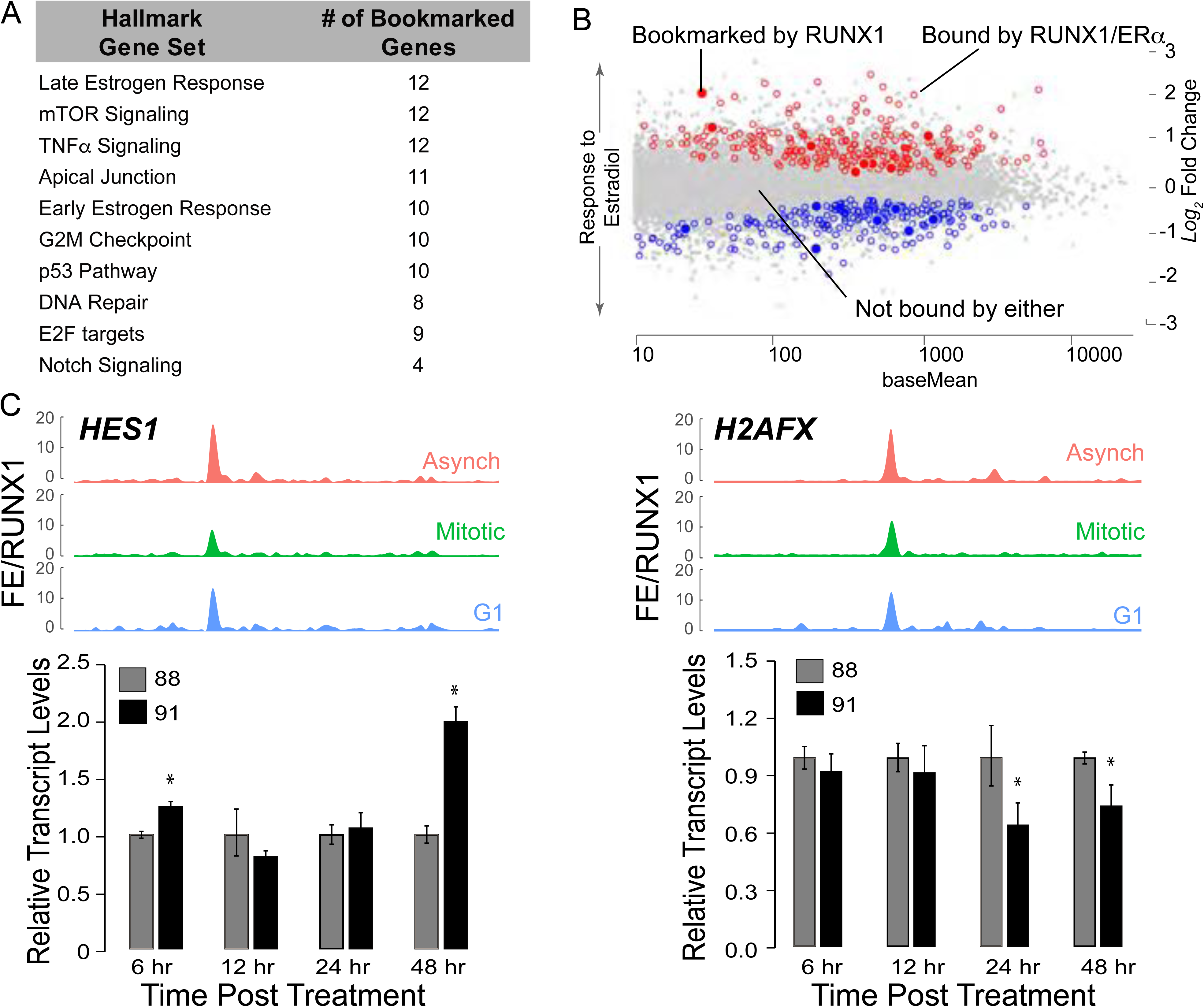
RUNX1 bookmarks RNA Pol II-transcribed genes involved in maintenance of breast epithelial phenotype. A) Gene Set Enrichment (GSE) analysis from interrogating mitotically bookmarked genes (i.e. RUNX1 mitotically occupied) against Hallmark Gene sets from Molecular Signatures Database (MSigDB). The top 10 most significantly overlapping gene sets are shown from top to bottom. B) Scatter plot of genes identified to be up or down regulated in response to estradiol treatment, that are also bound by estrogen receptor α (ERα) and RUNX1 (empty circles, blue for downregulated and red for upregulated). Scatter plot also illustrates up or down regulated genes in response to estradiol treatment that are bound by ERα and mitotically bookmarked by RUNX1 (filled in circles, blue for downregulated and red for upregulated). C) Top panel: ChIP-Seq tracks of *HES1* (left) and *H2AFX* (right) from asynchronous (top-red), mitotic (middle-green), and G1 (bottom-blue). Bottom panel: qRT-PCR data of *HES1* (left) and *H2AFX* (right) in asynchronous MCF10A cells treated with either active (AI-14-91) or inactive (AI-4-88) inhibitors for 6hr, 12hr, 24hr and 48hr at 20μM. Expression of target genes were normalized relative to beta actin.

We identified a novel subset of genes that are bookmarked by RUNX1 and relate to regulatory pathways involved in cellular phenotype including TNFα, Apical Junction and Notch signaling (Fig. 4A). Furthermore, NEAT1 and MALAT1, lncRNAs often deregulated in breast cancer (91, 92), were also mitotically bookmarked by RUNX1. Of the 413 RUNX1 bookmarked protein coding genes, *TOP2A*, *MYC*, *HES1*, *RRAS*, *H2AFX*, and *CCND3* are representative of RNA Pol II-transcribed genes involved in phenotype maintenance and cell fate decisions (See Suppl. Table 1 for complete list). Recently, *HES1* and *H2AFX* have been identified as regulators of breast epithelial phenotype (93–95). In our ChIP-seq dataset, *HES1* and *H2AFX* show significant fold enrichment of RUNX1 occupancy between the three populations of MCF10A cells (Fig. 4C; top panels). Expression of *HES1* increased upon inhibition of RUNX1 DNA binding (Fig. 4C; left panel—bar graph), indicating that RUNX1 represses *HES1*. In contrast, *H2AFX* expression at 24hr and 48hr of inhibitor treatment was decreased, suggesting RUNX1 activates H2AFX expression (Fig. 4C; right panel—bar graph). These results suggest that RUNX1 bookmarks both protein coding and non-coding genes that are critical determinants of epithelial lineage identity as a potential mechanism to stabilize the mammary epithelial phenotype.

### Inhibition of RUNX1 DNA binding causes epithelial to mesenchymal transition and alters the associated transcriptome

To experimentally address whether disruption of RUNX1 bookmarking leads to a change in epithelial phenotype, we treated cells with RUNX1 DNA binding inhibitor and the inactive control compound and monitored changes in cell morphology (Fig. 5A). Consistent with RUNX1 bookmarking and regulation of genes critical for epithelial phenotype (Figs. 2 & 4), disruption of RUNX1 DNA binding for 48 hours resulted in mesenchymal morphology. We next examined whether long-term inhibition of RUNX1 caused a permanent change in cell phenotype. Longer term treatment (5 days) of actively proliferating MCF10A cells showed significant apoptosis, although a small sub-population of cells survived and exhibited an altered phenotype (Fig. 5B). The surviving sub-population at day 5 was recovered by culturing cells in media without the inhibitor. By day 3-4 following media replacement, cells clearly showed a mesenchymal morphology (Fig. 5B), indicating that interfering with RUNX1 DNA binding causes loss of the normal mammary epithelial phenotype. Consistent with changes in cell morphology, we find alterations in expression and localization of the cytoskeletal F-actin protein (Fig. 5C). These observations were confirmed by examining the expression of epithelial markers (e.g., E-cadherin (Fig. 5D)), as well as mesenchymal markers (e.g., Vimentin (Fig. 5D)). E-cadherin was largely unchanged; however, Vimentin expression was significantly increased, confirming an epithelial-to-mesenchymal transition upon inhibition of the RUNX1-CBFβ interaction.

**Figure 5.**
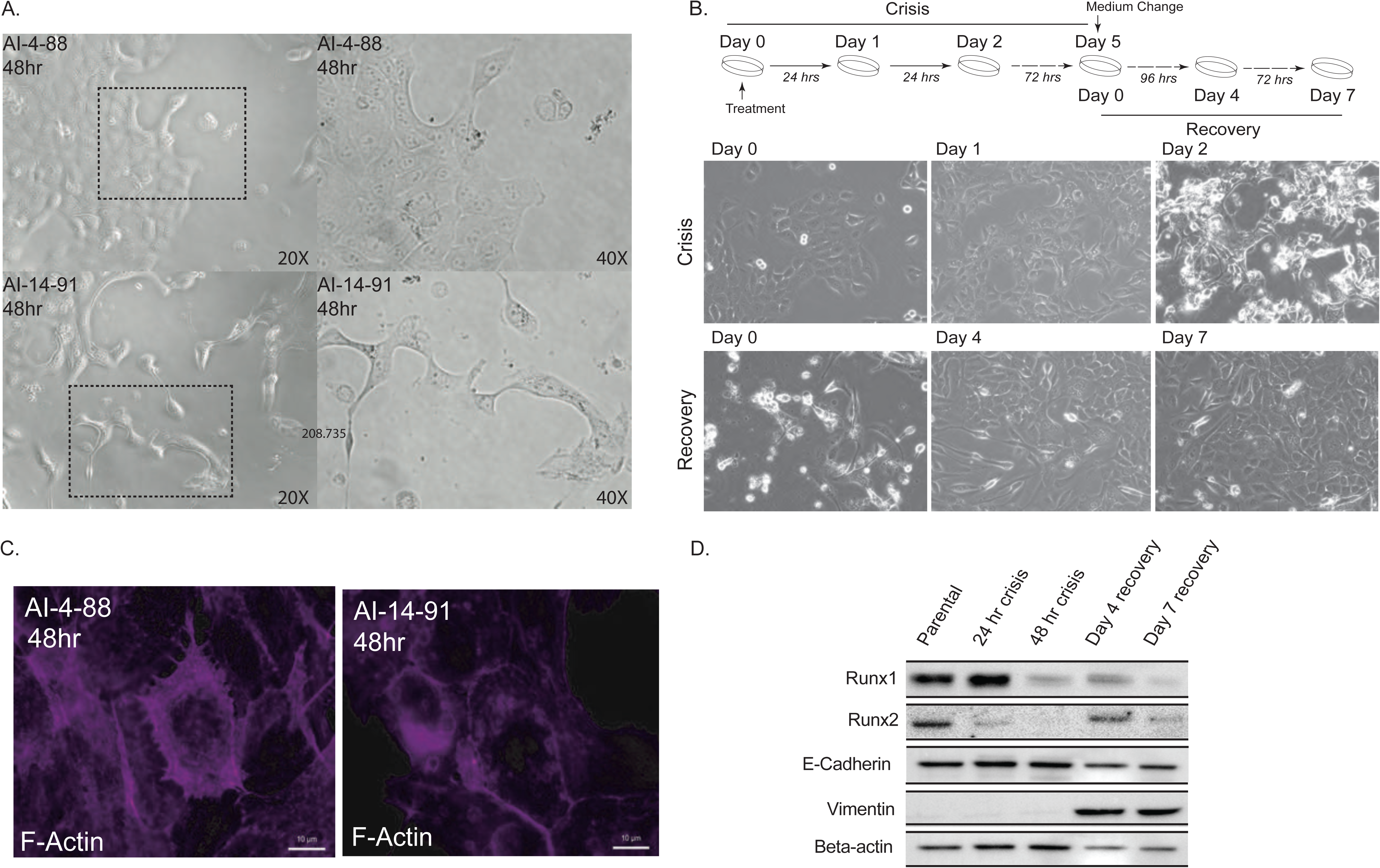
Disrupting RUNX1 mitotic gene bookmarking in MCF10A cells leads to a transformed cellular phenotype and EMT. A) Phase contrast microscopy images of MCF10A cells treated with AI-4-88 or A-14-91 for 48 hours at 20µM. Left panel – 20X magnification, right panel – 40X magnification. The outlines rectangle in the left panel is the resulting 40X magnification in the right panel. B) Top panel: Experimental schematic depicting treatment schedule for the “crisis” and “recovery” stages. Bottom panel: Phase contrast microscopy images from Day 0, 1, and 2 or crisis where MCF10A cells were treated with AI-14-91 at 20µM (top – left, middle, right respectively). Phase contrast images from Day 0, 4, and 7 or recovery following a media replacement. C) Morphological changes upon inhibition of the cytoskeletal protein F-actin. When compared to inactive compound (left panel), cells treated with the active compound show substantial alteration in cytoarchitecture (right panel). D) Western blot for RUNX1, RUNX2, epithelial marker E-cadherin, mesenchymal marker Vimentin, and loading control Beta-actin (top panel to bottom panel respectively) in MCF10A whole cell lysate harvested from the crisis 24 hour and 48-hour timepoints and recovery day 4 and day 7 timepoints.

To identify transcriptome-wide changes associated with EMT upon inhibition of RUNX1 DNA binding activity that is required for retention of the protein with mitotic chromosomes (56), we performed RNA sequencing (in triplicates) at each of the indicated time points (Day 1 and 2 in Crisis Phase and Day 4 and 7 in Recovery Phase). Heatmap analysis of all time points identified substantial changes in gene expression as cells transition from epithelial to mesenchymal phenotype (Fig. 6A). A differential gene expression analysis between the crisis and recovery phases uncovered significant changes in expression of genes associated with EMT (e.g., IL32, SERPINB2 etc.; Fig. 6B). Importantly, a subset of differentially expressed genes was occupied by RUNX1 during mitosis. We performed pathway analysis on differentially expressed genes (Fig. 6C and Suppl. Table 2). Consistent with phenotypic changes, we found that multiple signaling pathways that include TNF alpha, Interferon Gamma and estrogen responsiveness were altered during EMT caused by RUNX1 inhibition.

**Figure 6.**
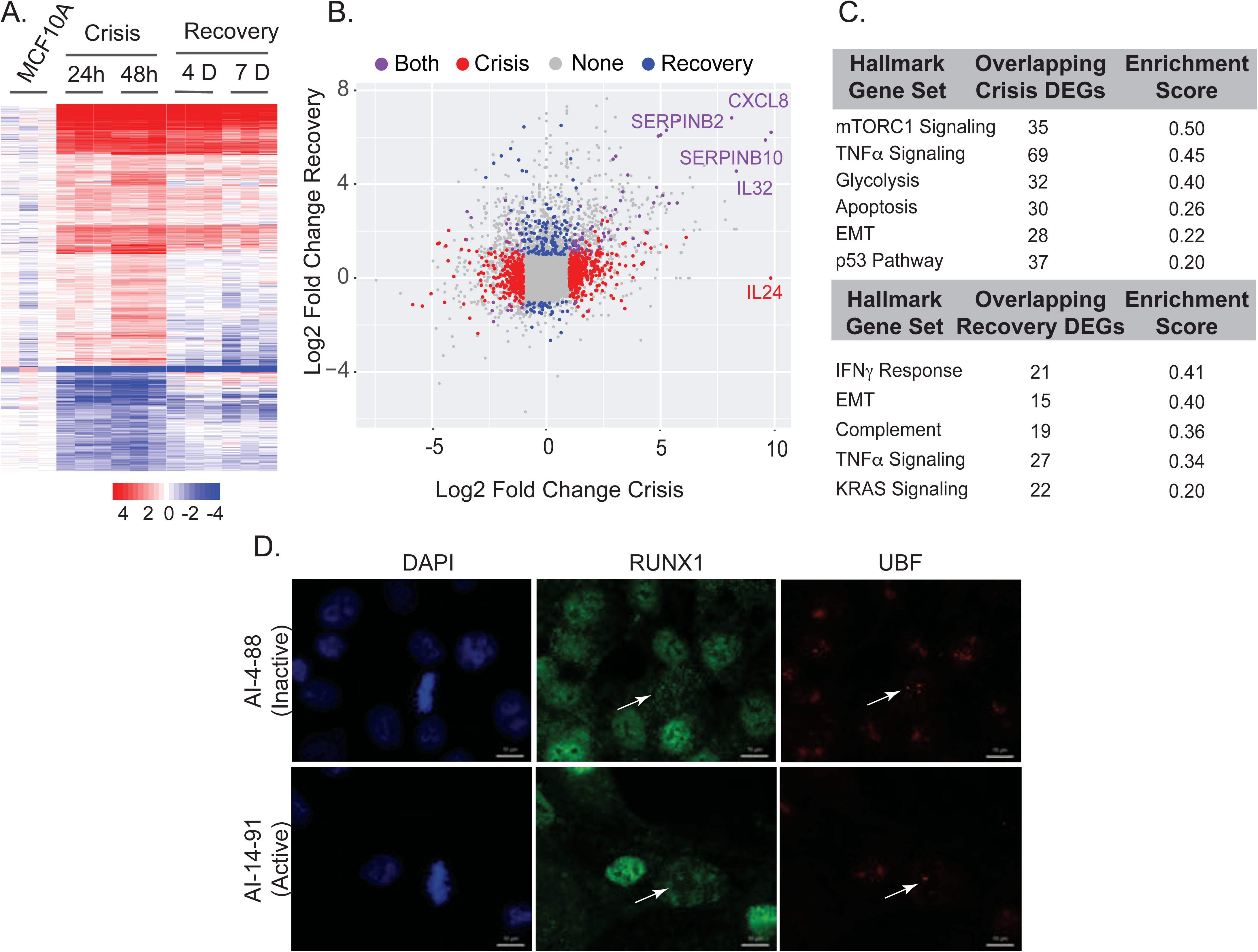
Differential expression and pathway analysis of RNA-Seq shows changes in key regulatory pathways involved in cell proliferation, metabolism, cell cycle control, estrogen response, and EMT. A) Expression heatmap of three biological replicates of 24-hour and 48-hour crisis timepoints and Day 4 and Day 7 recovery timepoints. B) Scatterplot of log2 fold change between crisis and recovery phases. Most changed genes in each stage are indicated. C) Table of overlapping pathways specific to crisis and recovery stages. D) Representative immunofluorescence images of the active compound (AI-14-91)-treated MCF10A cells in prophase and metaphase are shown. A substantial decrease in smaller RUNX1 foci (green) during mitosis is observed when compared to the inactive (AI-4-88) compound. Large RUNX1 foci that colocalize with UBF (red) are detectable at all substages of mitosis (white arrows) in the presence of either active or inactive compounds.

We next determined the effect of the inhibition of RUNX1-CBFβ interaction on mitotic retention of the protein. Actively proliferating MCF10A cells were treated with the inhibitor (AI-14-91) for 6hr, 12hr, 24hr, and 48hr at 20μM; a structurally similar but inactive compound (AI-4-88) was used as a control under identical conditions. Cells were subjected to immunofluorescence microscopy followed by detection of RUNX1 and UBF as described above. Although RUNX1 signal was detected in all mitotic sub-stages (Suppl. Fig. 5), we observed a substantial decrease in RUNX1 signal intensity on metaphase chromosomes (white arrows; Fig. 6D), indicating that RUNX1-CBFβ interaction and RUNX1 DNA binding activity play a key role in mitotic gene bookmarking. These changes were more pronounced for smaller RUNX1 foci and were not observed in control-treated cells; appreciable signal for large RUNX1 foci that colocalize with UBF (Fig. 3B & Suppl. Fig. 4) remained detectable in all sub-stages of mitosis (Fig. 6D and Suppl. Fig. 5). Together, these findings indicate that RUNX1 mitotic bookmarking of genes related to epithelial cell growth, proliferation, and lineage is associated with epithelial cell identity, and disruption of RUNX1 DNA binding leads to mesenchymal transition.

## DISCUSSION

This study identifies retention of RUNX1 on mitotic chromosomes and occupancy of target genes as a potential epigenetic mechanism for coordinate regulation of RNA Pol I- and II-transcribed genes that are critical for mammary epithelial proliferation, growth, and phenotype maintenance. Pharmacological inhibition of RUNX1 DNA binding causes transition of mammary epithelial cells to a mesenchymal phenotype, indicating that RUNX1 bookmarking of target genes contributes to stabilizing the normal breast epithelial phenotype.

Our findings are the first to examine RUNX1 bookmarking of target genes during mitosis in mammary epithelial cells and to report that RUNX1 coordinately controls cell growth-related ribosomal RNA (rRNA) genes and a large subset of cell proliferation/phenotype-related genes in these cells. One target gene of interest is hairy and enhancer of split-1 (*HES1*). Hes1 is a transcription factor that represses genes involved in cellular development and is regulated primarily by NOTCH signaling, one of our top ten overlapping hallmark gene sets bookmarked by RUNX1 (Fig. 4)(96, 97). HES1 was recently shown to have a prominent role in proliferation and invasion of breast cancer cells, and its silencing led to downregulation of p-Akt signaling and ultimately prevented EMT (93). Our findings indicate that RUNX1 stabilizes the normal mammary epithelial phenotype, in part, by bookmarking *HES1* and suppressing its expression.

Another important RNA Pol II-transcribed gene mitotically bookmarked by RUNX1 and critical for maintaining cellular phenotype is histone variant H2AFX (*H2AFX*). Silencing *H2AFX* in breast epithelial cells leads to induction of EMT through activation of *SNAIL2/SLUG* and *TWIST1* (95). Upon inhibition of the RUNX1-CBF*β* interaction, we find a decrease in *H2AFX* expression and a concomitant, significant increase in SNAIL2/SLUG expression. These data identify RUNX1 as a novel upstream regulator of *H2AFX* expression; RUNX1 bookmarking and activation of H2AFX and subsequent suppression of *SNAIL2/SLUG* prevents EMT in breast epithelial cells.

Several groups have shown that RUNX1 interacts with ERα at both enhancer regions and transcriptional start sites (TSSs) for regulation of specific genes (34, 88). Our ChIP-Seq results, coupled with publicly available data sets, reveal a novel finding: RUNX1 bookmarking of a subset of ERα-occupied, hormone-responsive genes during mitosis may be critical for maintenance of the breast epithelial phenotype. Future studies will be required to investigate mechanistic significance of this observation in estrogen receptor positive mammary epithelial and breast cancer cells.

Our findings suggest that mitotic gene bookmarking by RUNX1 contributes to regulation of the mammary epithelial phenotype. Equally important, our study shows that inhibition of RUNX1 DNA binding specifically elicits an epithelial-to-mesenchymal transition that accompanies changes in critical genes and pathways involved in EMT. These findings are further supported by RUNX1 mitotic occupancy of cell growth-related rRNA genes, and together highlight a key role of RUNX1 in coordinating cell proliferation, growth and phenotype. Because RUNX1 interacts with multiple co-activators and co-repressors, additional in-depth studies are required to determine contributions of RUNX1 co-regulatory proteins to mitotic gene bookmarking.

Another novel contribution of the current study is mitotic bookmarking of lncRNAs by a transcription factor. RUNX1 was recently shown to regulate lncRNAs NEAT1 and MALAT1 (89, 91), which have critical roles in the onset and progression of breast cancer (92). Our findings show that, in addition to occupying protein coding genes, RUNX1 bookmarks several lncRNAs for post-mitotic regulation. It will be important to determine if RUNX1-bookmarked lncRNAs have G1-specific roles in maintaining the normal mammary epithelial phenotype and/or in the onset and progression of breast cancer.

In summary, this study shows that RUNX1 occupies RNA Pol I- and II-transcribed genes during mitosis for coordinate regulation of normal mammary epithelial proliferation, growth, and phenotype. Disruption of RUNX1 DNA binding leads to the epithelial-to-mesenchymal transition, a key event in breast cancer onset, and implicates RUNX1 mitotic gene bookmarking as an epigenetic mechanism to physiologically sustain the mammary epithelial phenotype.

## MATERIALS AND METHODS

### Cell Culture Techniques

Breast epithelial (MCF10A) cells were cultured in DMEM/F-12 50/50 mixture (Corning™, Corning, NY). Culturing media was also supplemented with horse serum to 5% (GIBCO, Grand Island, NY), human insulin to 10µg/mL (Sigma Aldrich, St. Louis, MO), human epidermal growth factor to 20ng/mL (PeproTech, Rocky Hill, NJ), cholera toxin to 100ng/mL (Thomas Scientific, Swedesboro, NJ), hydrocortisone to 500ng/mL (Sigma Aldrich, St. Louis, MO), Penicillin-Streptomycin to 100U/mL (Thermo Fisher Scientific, Ashville, NC), and L-Glutamine to 2mM (Thermo Fisher Scientific, Ashville, NC).

For mitotic arrest of parental MCF10A cells, culturing media was supplemented with 50ng/mL of Nocodazole (Sigma Aldrich, St. Louis, MO) and incubated with cells for 16hrs. Supplementing culturing media with equivalent volumes of DMSO (Sigma Aldrich, St. Louis, MO) served as a control. DMSO-treated and mitotically arrested populations of MCF10A cells were harvested following the 16hr incubation. For G1 (released from mitotic arrest) populations of MCF10A cells, the nocodazole-supplemented media was replaced with normal media and cells were incubated for 3hrs, after which the released cell population was harvested for subsequent analyses that include western blotting, qPCR, FACS and ChIP-seq.

### Western Blot Analyses

Protein lysates were prepared by incubating cells in RIPA buffer on ice for 30min, followed by sonication using Q700 Sonicator (QSonica, Newtown, CT). Proteins were resolved by SDS-PAGE and transferred to PVDF membrane using standard protocols. Following primary antibodies were used at a 1:1000 dilution (except Lamin B, which was used at 1:2000 dilution) in this study: UBF (sc-13125, Santa Cruz Biotechnology, Dallas, TX); RUNX1 (4334S, Cell Signaling Technologies, Danvers, MA); Cyclin B (4138S, Cell Signaling Technologies, Danvers, MA); Beta-Actin (3700S, Cell Signaling Technologies, Danvers, MA), and CDT1 (ab70829, AbCam, Cambridge, UK); Lamin B1 (ab16048, AbCam, Cambridge, UK). Horseradish peroxidase conjugated secondary antibodies used in this studies were: goat anti-mouse IgG at 1:5000 dilution (31460, Invitrogen, Carlsbad, CA), goat anti-rabbit IgG HRP conjugated (31430, Thermo Fisher Scientific, Ashville, NC) at 1:1000, 1:2000, or 1:5000 dilutions. Blots were developed using Clarity Western ECL Substrate (Bio-Rad, Hercules, CA) and imaged using Molecular Imager^®^ Chemi doc™ XRS+ Imaging System (Bio-Rad, Hercules, CA) aided by Image Lab Software Version 5.1 (Bio-Rad, Hercules, CA).

### Confocal Microscopy, Image Acquisition, Processing and Analyses

MCF10A cells were plated on gelatin-coated coverslips in 6-well plates at 175,000 cells/mL and processed for immunofluorescence 24 hrs after plating using standard protocol. Briefly, cells were washed twice with sterile-filtered PBS on ice and cell were fixed in 1% MeOH-free Formaldehyde in PBS for 10min. After permeabilization in 0.25% Triton X-100-PBS solution, cells were sequentially incubated with primary and Alexa fluorophore conjugated secondary antibodies for 1hr each at 37°C in a humidified chamber with extensive washes after each incubation. Primary antibodies used were: RUNX1 at 1:10 dilution (4334S, Cell Signaling Technologies, Danvers, MA), and Upstream Binding Transcription Factor (UBF) at 1:200 dilution (F-9 sc-13125, Santa Cruz Biotechnology, Dallas, TX). Secondary antibodies used were goat anti-rabbit IgG conjugated with Alexa Fluor 488 (A-11070, Life Technologies, Carlsbad, CA) and goat anti-mouse IgG conjugated with Alexa Fluor 594 (A-11005, Life Technologies, Carlsbad, CA) diluted 1:800. Cells were counterstained with DAPI to visualize DNA and coverslips were mounted onto slides in ProLong Gold Antifade Mountant (Thermo Fisher Scientific, Ashville, NC). Images were captured using a Zeiss Axio Imager.Z2 fluorescent microscope and Hamamatsu ORCA-R2 C10600 digital camera. Images were processed using ZEN 2012 software.

To examine mitotic localization of RUNX1 in unfixed cells, an expression plasmid containing RUNX1-EGFP was introduced using either nucleofection or Lipofectamine 2000 transfection reagent in actively proliferating MCF10A cells grown on gelatin-coated coverslips. After 16 hours of nucleofection, cells were washed once with 1X PBS and stained with Hoechst dye to visualized DNA. Coverslips were mounted using the ProLong Gold Antifade Mountant and subjected to confocal microscopy.

Cells were initially imaged with a Zeiss LSM 510 META confocal laser scanning microscope (Carl Zeiss Microscopy, LLC., Thornwood, NY, USA). The DAPI signal was excited with a 405 nm laser and collected with a 425-475 nm band pass filter, Alexa 488 was excited with a 488 nm laser and collected with a 500-550 nm band pass filter, and Alexa 568 with a 561 nm laser and collected with a 570-620 nm band pass filter. Images were captured with a Plan-Fluor 40X (1.3 NA) objective lens. The confocal pinhole was initially set to 1.2 Airy Unit diameter for the 561 nm excitation giving an optical section thickness of 0.41 μm. Images were acquired at 12-bit data depth, and all settings, including laser power, amplifier gain, and amplifier offset were established using a look up table to provide an optimal gray-scale intensities. All images were acquired using matching imaging parameters. The acquired images were subjected to colocalization analysis via Volocity version 6.3.0 (Perkin Elmer, Waltham, MA, USA). At least 15 interphase and 15 metaphase cells were identified in captured images and appropriate thresholds were manually determined to eliminate background fluorescence for calculating Pearsons and Manders correlation coefficients between RUNX1 and UBF.

To ensure the specificity of our observations, additional samples were imaged with a Nikon A1R-ER laser scanning confocal microscope (Nikon, Melville, NY, USA). Images were acquired with the galvano scanner at a frame size of 1024 × 1024 pixels with an Apo TIRF 60X objective lens (N.A. 1.49) zoom of 2 and 1.2 Airy Unit pinhole setting. Images were also viewed in NIS Elements version 5.02.01 and analyzed using the line profiling tool. Overlaying DAPI, RUNX1, and UBF fluorescent intensities from individual channels along the line profile revealed overlapping peak intensities between the RUNX1 and UBF channels, thus indicating colocalization.

### RNA Isolation, cDNA Synthesis and Quantitative PCR

Total RNA was isolated from MCF10A cells using TRIzol™ Reagent (Invitrogen, Carlsbad, CA) and Direct-Zol™ RNA MiniPrep isolation kit (Zymo Research, Irvine, CA) following manufacturer instructions. cDNA was created using SuperScript IV® First-Strand Synthesis System for RT-PCR (ThermoFisher, Asheville, NC). Resulting samples were quantified on a Qubit 2.0 Fluorometer (Invitrogen, Carlsbad, CA) and diluted to 500pg/μL. Equal amounts of DNA template were loaded for samples analyzed by qPCR. At least three independent biological replicates were analyzed for expression of RUNX1 bookmarked genes by qPCR. Student’s t-test was used to determine the significance of changes in transcript levels under different biological conditions.

### Chromatin Immunoprecipitation, Library Preparation, Sequencing and Bioinformatics Analyses

Asynchronous (Asynch), mitotically arrested (M), and released from mitosis (G1) MCF10A breast epithelial cells were subjected to chromatin immunoprecipitation using a modified Farnahm’ protocol {O’Geen, 2010 #2713}. Sonication parameters for each population of cells was as follows: Peak Watt 140W, Duty Factor 10, Cycle/Burst 200 using a S220 focused ultra-sonicator (Covaris, Matthews, NC). M and G1 populations of cells were sonicated for 28min total, whereas asynchronous populations of cells were sonicated for 36min. An aliquot of sonicated lysates was boiled in 100°C for 15min with NaCl and elution buffer and DNA was purified using PureLink™ PCR Purification Kit (K310001, ThermoFisher, Ashville, NC). Purified DNA was resolved on a 1.5% agarose gel to confirm optimal sonication (bulk of fragments between 200-400bp) prior to performing ChIP. In parallel, an aliquot was also quantified via Qubit 2.0 Fluorometer (Invitrogen, Carlsbad, CA) and analyzed by using a High Sensitivity DNA Kit on a Bioanalyzer 2100 (Agilent, Santa Clara, CA).

For chromatin immunoprecipitation (ChIP) reactions, 150ug of sonicated chromatin was incubated with 10ug of RUNX1 antibody (4336BF, Cell Signaling Technologies, Danvers, MA), diluted 1:10 in IP dilution buffer, and incubated overnight (16-18hrs) at 4°C with mild agitation. Following incubation, 150uL of Protein A/G magnetic beads (Thermo Scientific – Pierce, Waltham, MA) per ug of antibody used were added to each IP reaction and incubated for 2-4hrs at 4°C with mild agitation. Beads were extensively washed with IP wash buffers and resuspended in Elution Buffer to extract immunoprecipitated chromatin, which was subsequently purified using PureLink™ PCR Purification Kit. At least 3 biological replicates were performed for each cell population and each antibody.

ChIP libraries were generated using Accel-NGS^®^ 2S Plus DNA Library kit (Swift Biosciences, Ann Arbor, MI) following manufacturers protocol. Input and RUNX1 ChIP samples were normalized to 1ng prior to library generation. Libraries were amplified in an optional PCR step for 12 total cycles. Finalized libraries were double size selected using AMPure XP beads (0.8X and 0.2X volume ratios to sample), resulting in the majority fragments sized between 250-400bp. Next generation sequencing of pooled ChIP libraries was performed by the University of Vermont Cancer Center - Vermont Integrated Genomics Resource (VIGR).

Single end, 50bp reads (SE50) were processed pre-alignment by removing adapter reads (Cutadapt v1.6) and trimming low quality base calls from both ends (FASTQ Quality Trimmer 1.0.0; min score >= 20, window of 10, and step size of 1). Because we were specifically investigating rDNA, a customized build of hg38 was constructed that included normally masked regions of rDNA (Gencode U13369). Since some (although not complete) rDNA sequence is present in the hg38 assembly, we masked all parts of hg38 that would normally be attributed to rDNA sequences (bedtools v2.25.0 maskfasta) based on alignment positions of 50 bp *in silico* reads generated across U13369. Finally, we appended the complete rDNA sequence as a distinct sequence (chrU13369.1) to the masked hg38 FASTA resulting in the hg38_rDNA assembly used for analysis.

Resulting reads were aligned to hg38_rDNA (STAR v2.4; splicing disabled with ‘-- alignIntronMax 1’). Next, we called peaks and generated fold-enrichment (FE) bedGraph files (MACS2 v2.1.0.20140616; callpeak at p-value e-5; and bdgcmp with FE method)(98). Irreproducible Discovery Rate (IDR) was conducted using unpooled replicates with all peaks in pooled samples passing an IDR cutoff of 0.5 (99). To reduce artificial peaks, we calculated strand cross-correlation for all peaks at a shift of 95 bp (the mean observed fragment size of 180 bp minus the read size of 85bp) and unshifted (100). We eliminated peaks that exhibited low shifted correlation (shifted correlation <0.7) and those that exhibited high unshifted correlation relative to shifted (shifted – unshifted correlation < 0.1). This increased retrieval of the RUNX1 motif and improved agreement with other RUNX1 datasets. Passing peaks were annotated separately to mRNA and lncRNA transcript start sites (TSSs) using Gencode v27 with a distance cutoff of 5000 bp. Regional distribution of peaks was determined using the same annotation reference limited to the “basic” tag for exons and promoters.

### Inhibitor Treatment and Assessment of Global Protein Synthesis

Core binding factor – Beta (CBFβ) inhibitors AI-4-88 and AI-14-91 were kindly provided by John H. Bushweller (University of Virginia) and used to evaluate RUNX1 DNA-binding inhibition in MCF10A cells. Protein synthesis evaluation by immunofluorescence was conducted following manufacturer protocol (K715-100, BioVision, San Francisco, CA). To examine effects of inhibiting the RUNX1-CBFβ interaction, MCF10A cells were treated with active or inactive compound for 48 hours. Culture medium containing the active or inactive compounds was replaced with fresh medium without the compounds and cells were harvested 4- and 7-days post medium change.

### RNA-Sequencing, Differential Expression Analysis, and Pathway Analysis

RNA was isolated using Direct-zol RNA MiniPrep (Zymo Research, Irvine, CA, USA) and was quantified and assayed for RNA integrity by Bioanalyzer (Agilent Technologies, Inc., Santa Clara, CA, USA). Following the removal of ribosomal RNA, the RNA pool was reverse transcribed, amplified, purified, and bound to strand-specific adaptors following the manufacturer’s protocol (SMARTer Stranded Total RNA Sample Prep Kit, Takara Bio, Mountain View, CA, USA). cDNA libraries were assayed for quality control by Bioanalyzer (Agilent Technologies, Inc., Santa Clara, CA, USA). After cDNA quality validation, generated libraries were sequenced. 24 hour and 48-hour counts were grouped together into one “crisis” category and the day 4 recovery and day 7 recovery counts were grouped together into one “recovery” category. Treatment groups were compared with untreated MCF10A cells. After demultiplexing and quality filtering, reads were aligned to hg38 using Gencode (GRCh38.p13). As a reference, annotation with STAR (v2.5.2a)(101) aligned reads were then counted using HT-Seq (102). Differential gene expression was analyzed using DESeq2 in R v.3.5.1 (103). Parameters for significant differential expression were base mean expression greater than five, absolute log2 fold change greater than one, and a p-value less than 0.05. Pathway analysis was performed using Gene Set Enrichment Analysis v6.3 (Broad Institute, Inc., MIT, UC San Diego).

## ACKNOWLEDGEMENTS

The authors thank Dr. Prachi Ghule, Department of Biochemistry, University of Vermont for assistance in confocal microscopy, Scott Tighe, Pheobe Kehoe, and Jessica Hoffman of the Vermont Integrative Genomics Resource for performing next generation sequencing of samples, Dr. Roxana del Rio-Guerra of the Flow Cytometry and Cell Sorting Facility for analysis of samples by FACS and Dr. Alan Howe, Department of Pharmacology, University of Vermont, for providing Phalloidin reagent used in immunofluorescence microscopy experiments.

## DATA AVAILABILITY

GEO accession number for the sequencing data generated in this study is GSE121370.

## FIGURE LEGENDS

**Supplementary Figure 1.**
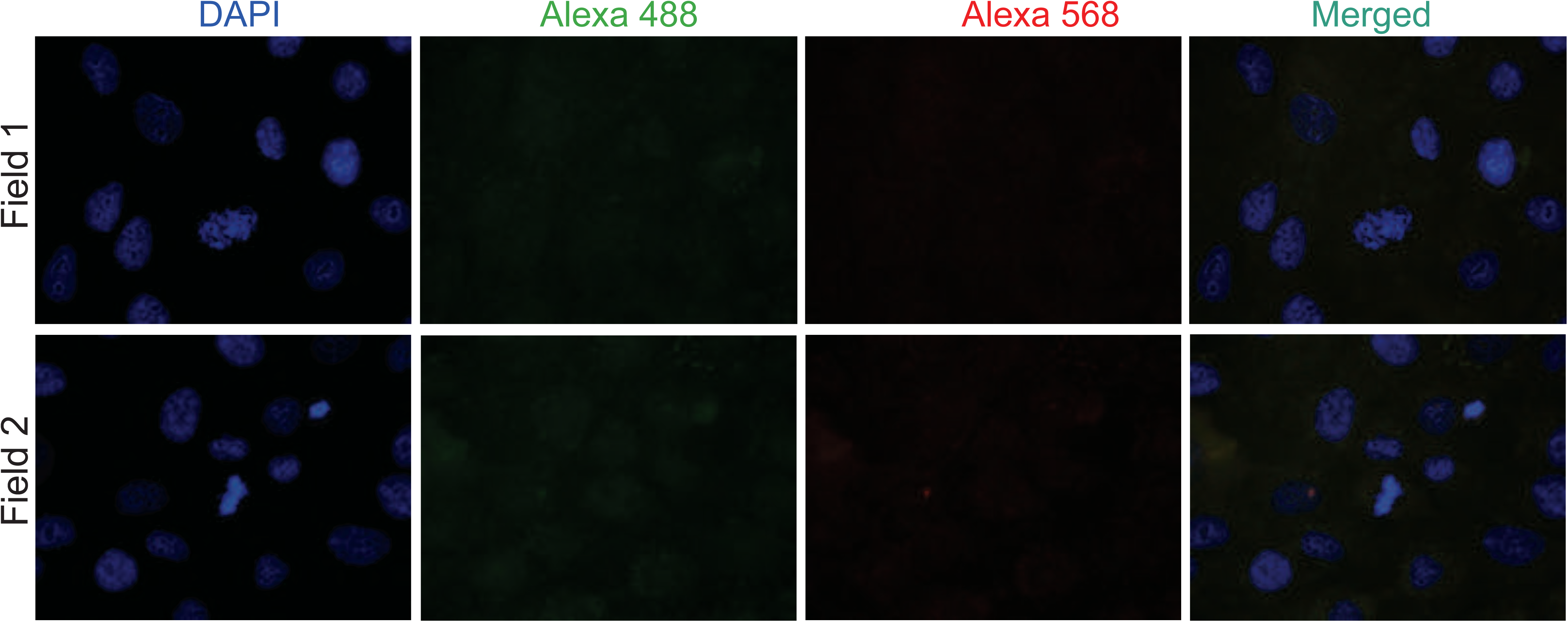
Secondary antibody controls for immunofluorescence microscopy. To ensure the specificity of RUNX1 signal on mitotic chromosomes, actively proliferating mammary epithelial MCF10A cells, grown on gelatin-coated coverslips, were subjected to immunofluorescence microscopy procedure as described in Materials and Methods section with one modification: no primary antibody was added, but secondary antibodies were used at the same dilution as in all IF experiments. Nuclei were counterstained with DAPI. Two different fields are shown, confirming that the RUNX1 signal we observe on mitotic chromosomes is specific.

**Supplementary Figure 2.**
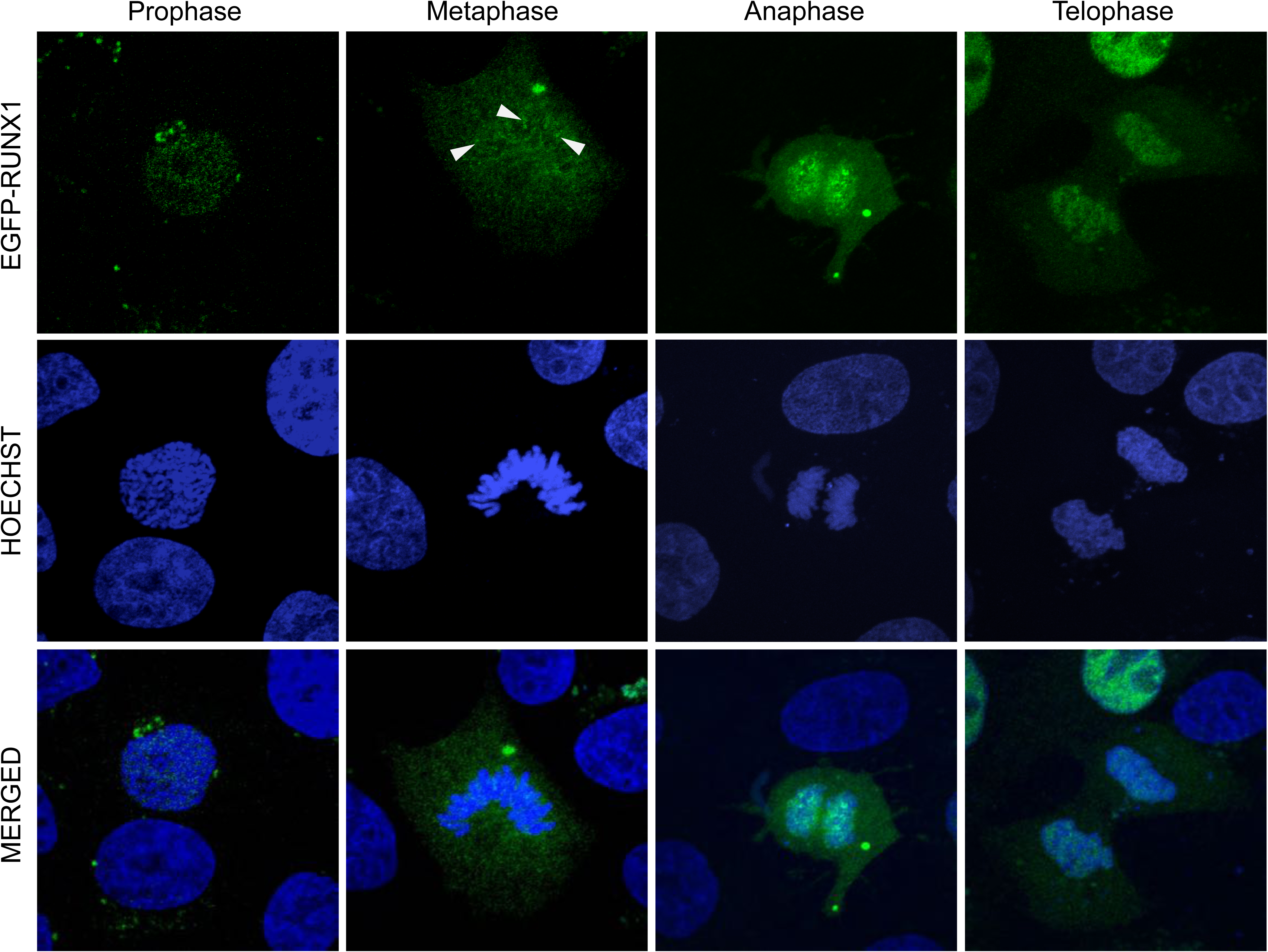
RUNX1 associates with chromosomes during all stages of mitosis in unfixed live MCF10A cells. Mammary epithelial MCF10A cells were transiently transfected with EGFP-RUNX1 and imaged by confocal microscopy without fixation (see Materials and Methods section for details). Top panels show RUNX1 (green) association with mitotic chromosomes in unfixed, live MCF10A cells. Cells were counterstained with Hoechst (middle panels; blue) to visualize DNA in live cells and to identify mitotic cells. Merged images (bottom panels) were generated to confirm localization of RUNX1 signal with DNA. Arrow heads indicate punctate RUNX1 foci retained on mitotic chromosomes.

**Supplementary Figure 3.**
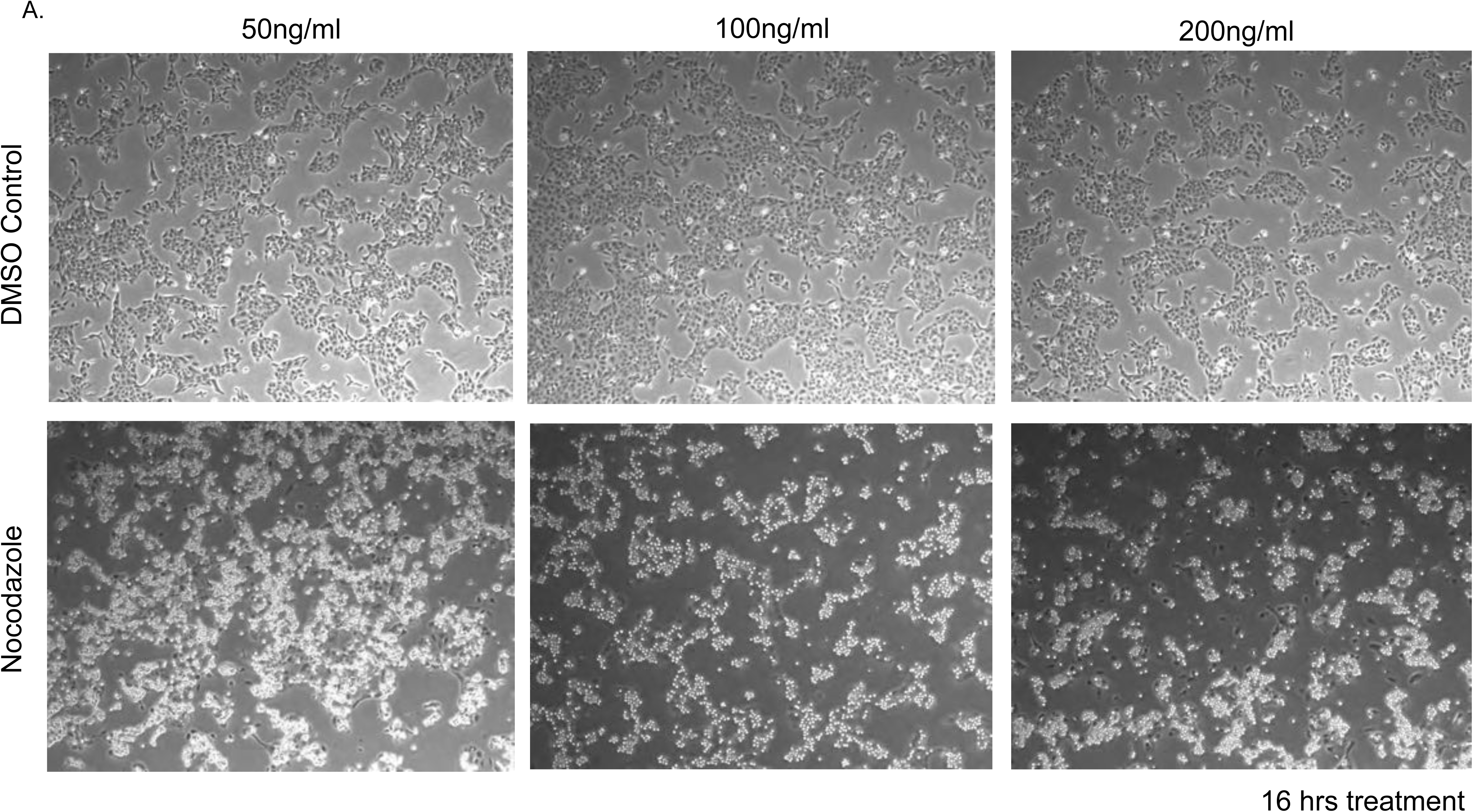
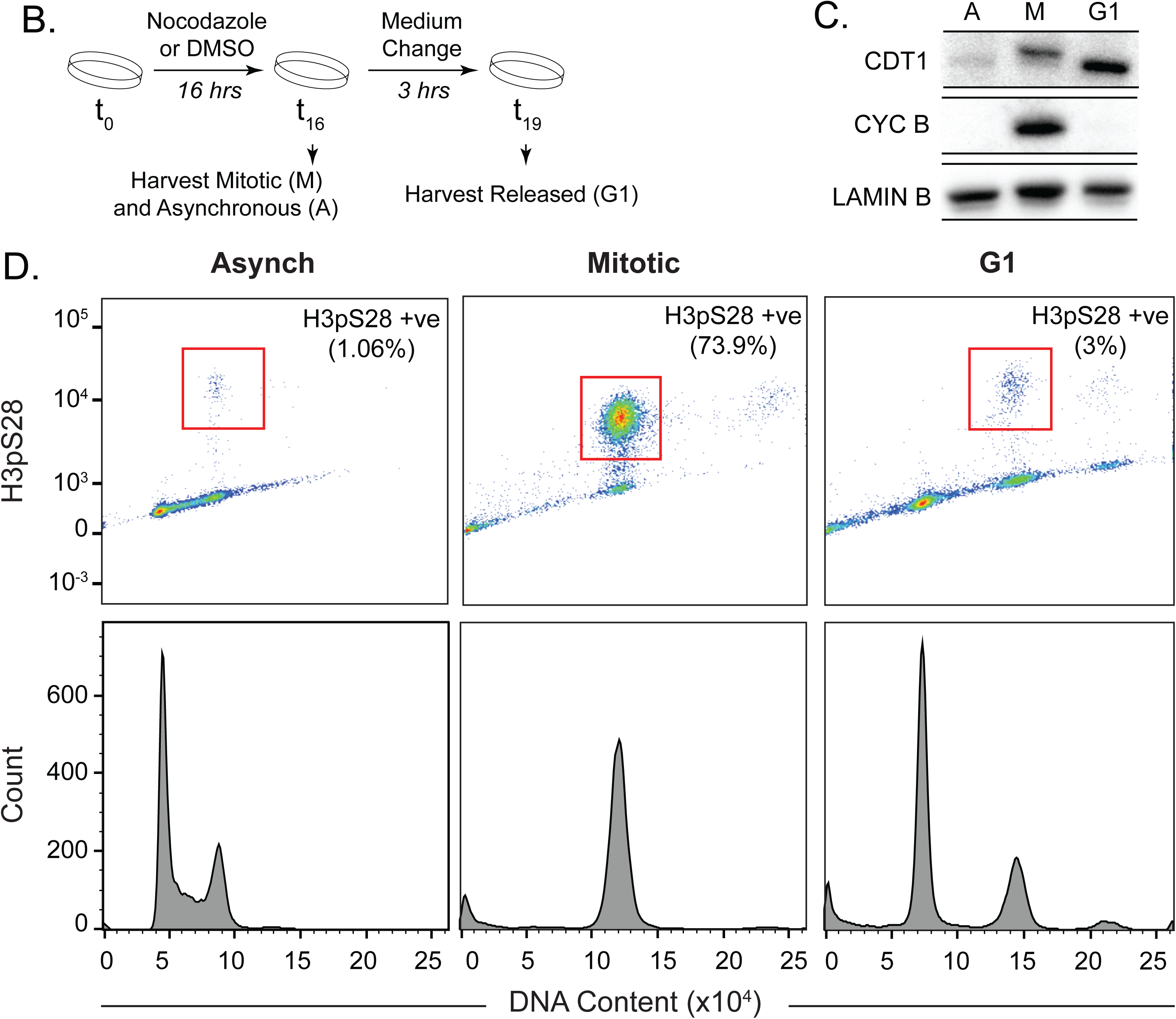
Determination of optimal nocodazole dosage for maximum mitotic block. A) Micrographs of MCF10A cells, treated with various doses of nocdazole for 16 hours, are shown (bottom panels). DMSO control treatment is also included (top panels). B) Experimental schematic depicting mitotic arrest and harvest of each treated MCF10A cell population: Asynchronous – A, Mitotic – M, and Released – G1. C) Western blot of each harvested MCF10A population for cell cycle specific markers to evaluate mitotic arrest and synchronization procedure. D) Fluorescence-activated cell sorting (FACS) analysis of harvested A, M, and G1 MCF10A populations to determine mitotic purity and DNA content (n=2 biological replicates per group).

**Supplementary Figure 4.**
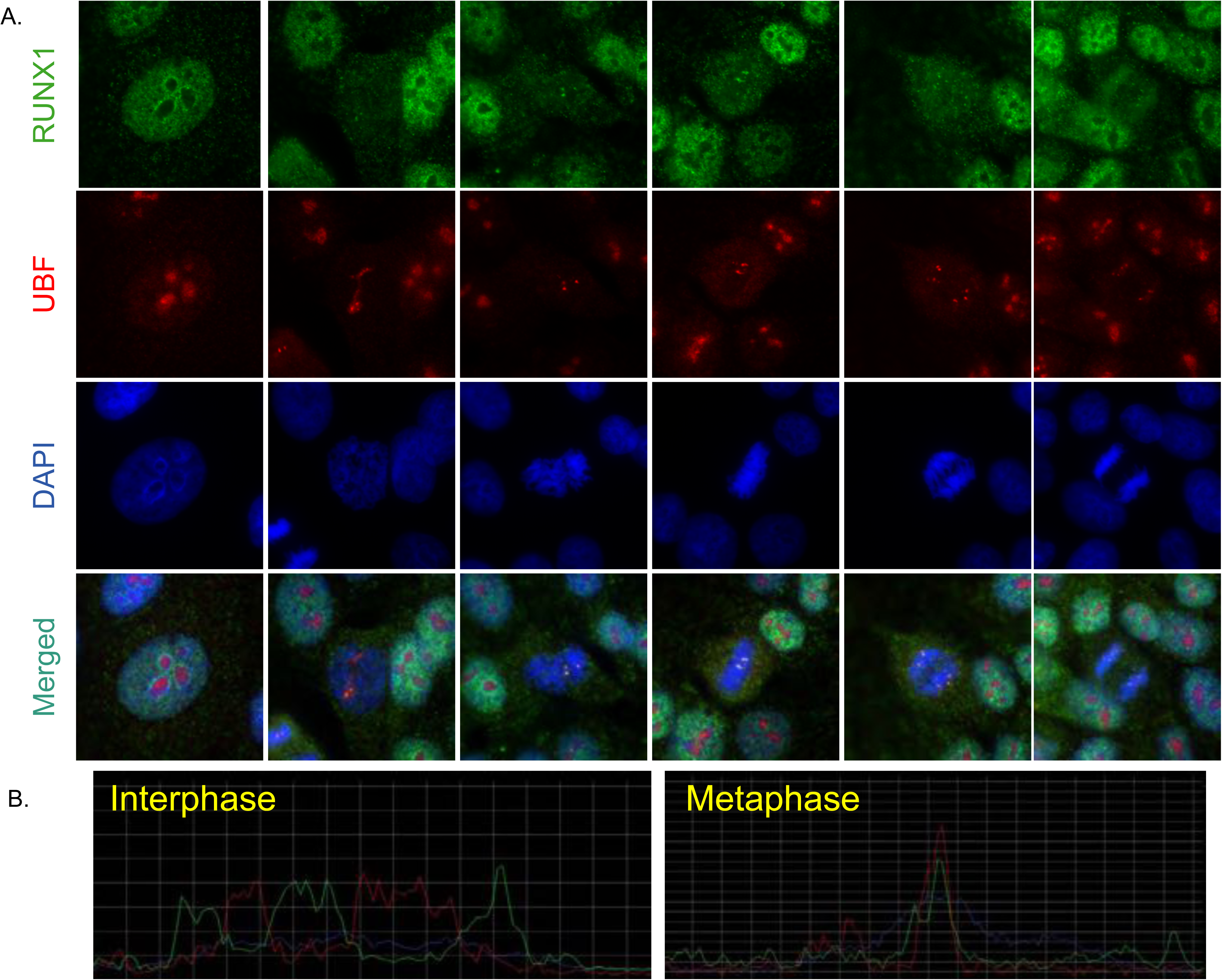
RUNX1 colocalizes with RNA Pol I subunit, upstream binding factor (UBF) on mitotic chromatin. A) Immunofluorescence microscopy images of RUNX1 (green – top row), UBF (red – 2^nd^ row from top), DAPI (blue – 2^nd^ row from bottom), and the three channels merged (bottom row) in MCF10A cells. Images were captured of spontaneously dividing MCF10A cells in different substages of mitosis. B) Representative images of line profiles taken on interphase vs metaphase cells (n=15 each).

**Supplementary Figure 5.**
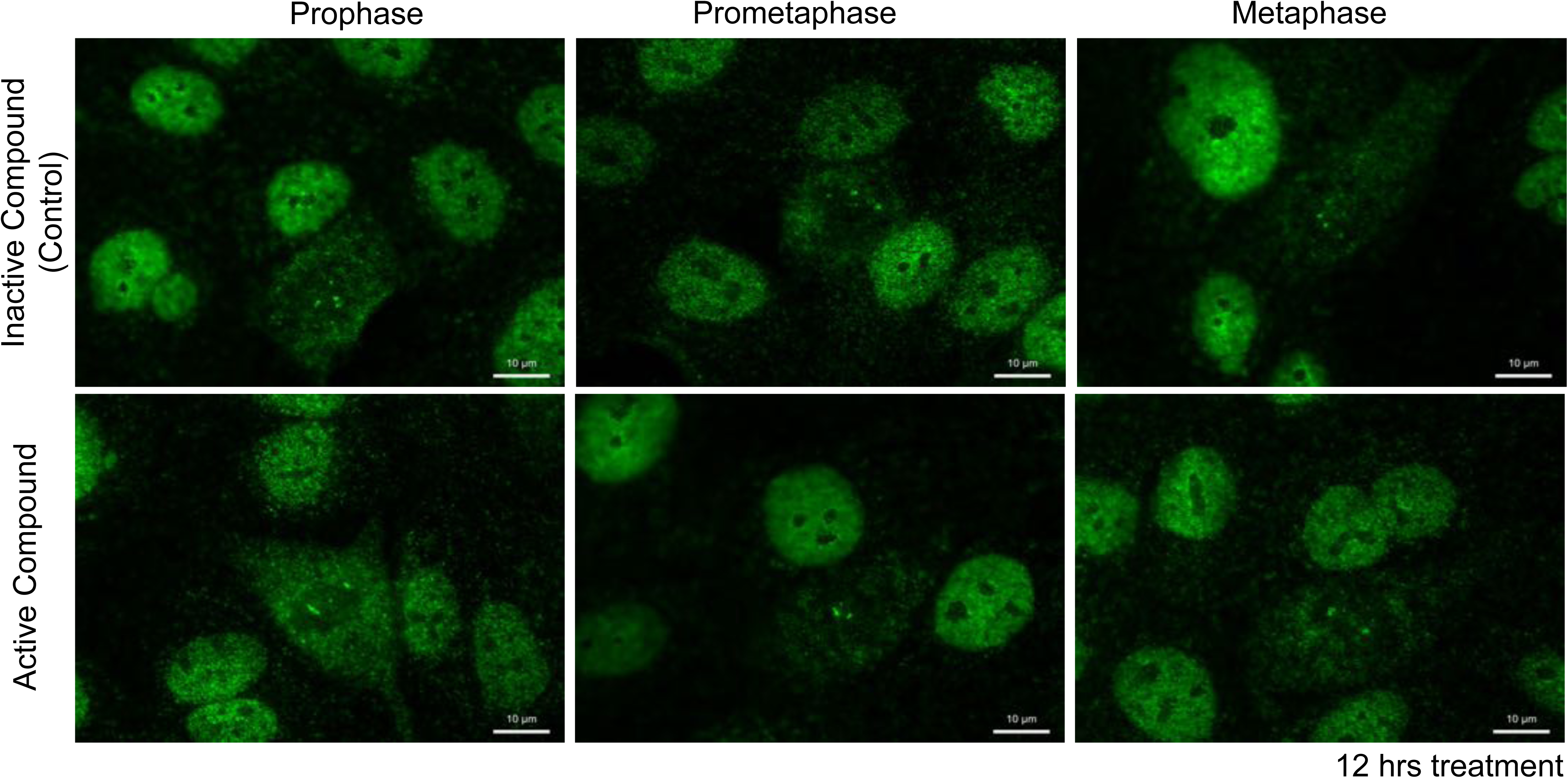
The RUNX1-CBFβ inhibitor reduces RUNX1 association with mitotic chromosomes. MCF10A cells, treated with 20µM inactive control compound or active inhibitor for 12 hours, were stained for localization of endogenous RUNX1 (green) to mitotic chromosomes. RUNX1 retention on mitotic chromosomes, particularly in smaller foci, was substantially reduced in cells treated with active inhibitor of RUNX1-CBFβ interaction, which disrupts RUNX1 DNA binding activity.

**Dataset S1 (separate file).** List of genes occupied by RUNX1 in asynchronous, mitotic and G1 cell populations in mammary epithelial cells.

**Dataset S2 (separate file).** Gene enrichment analysis of genes sensitive to RUNX1-CBFβ inhibition during crisis and recovery phases of epithelial to mesenchymal transition in MCF10A cells.

